# Evidence for recruitment-mediated decline in an Eastern box turtle (*Terrapene carolina carolina*) population based on a 30-year capture-recapture data set from Maryland

**DOI:** 10.1101/2024.08.28.610102

**Authors:** J. Andrew Royle, Michael M. Quinlan, Christopher W. Swarth

## Abstract

The Eastern box turtle (*Terrapene carolina carolina*) population at the Jug Bay Wetlands Sanctuary, Lothian, MD has been monitored continuously for 29 years (1995-2023). We used open population capture-recapture models (Jolly-Seber) to estimate annual population size, survival probability, and recruitment rate. The model allows for unknown sex of individuals and includes information on individuals found dead. Our analysis documents a long-term decline of approximately 67% in box turtle population size at the Sanctuary over this nearly three-decade period. We estimate annual survival for both males and females, which does not show a systematic increase or decrease over time, averaging about 0.90 (95% CI: 0.86, 0.93) for females and 0.97 (95% CI: 0.94, 0.98) for males. Conversely, per-capita recruitment shows a marked decline over the first 15 years of the record, suggesting that population declines may be due to reduced recruitment. Conservation efforts for the species could benefit from a formal population viability analysis to understand the relative effects of survival and recruitment on changes in population size for this long-lived species.

---

The Eastern box turtle (*Terrapene carolina carolina*) is a charismatic species that is widespread in eastern North America, even in the fragmented habitat matrix surrounding large cities. Although perceived to be relatively common (Roberts et al. 2024), it is considered a species of concern or species of greatest conservation need (SGCN) throughout its range (Erb and Roberts 2023), and the species is listed as vulnerable by the IUCN (van Dijk 2011). There has been increasing attention devoted to monitoring box turtle populations (Erb et al. 2015; Roberts and Erb 2023) and understanding causes and mechanisms of population declines (Nazdrowicz et al. 2008; Jones et al. 2021; Roberts et al. 2024).

Although some studies suggest substantial population declines over recent decades (Hall et al. 1999, Kemp et al. 2022), there are few contemporary studies that document long-term trends in Eastern box turtle populations. Kemp et al. (2022) is the most recent in the literature. They reported a decrease in population size of almost 75% over a 42-year period for a population from southeast Pennsylvania. However, their data were sparse over most of the period and this necessitated a relatively coarse-grained estimation of population size over two time periods, the early (1978-1982) and later (2015-2022) parts of the record. Hall et al. (1999) documented significant long-term declines of about 80% in the number of individuals encountered in an intensively studied population (Stickel 1950) over a 50-year period ending with a survey conducted in 1995. Their analysis did not use capture-recapture models and they did not report population size or survival estimates. Nazdrowicz et al. (2008) reported a decline of about 76% (from 91 individuals to 22) in a small Delaware population over the period 1968 to 2002. Conversely, a recent study from North Carolina (Roe et al. 2021) found “no evidence of population decline at any site over a ten-year period (2008–2017)” across small, local populations, based on an analysis of mostly opportunistic data from a large number of sites where sampling protocols and monitoring efforts differed. Because the site-specific data sets upon which their analysis was based are short-term, small and heterogeneous, the analysis likely had very low power to detect trends of any magnitude. Dodd et al. (2012) analyzed a 16-year study of the Florida box turtle (*T. carolina bauri*), showing a 5% per year increase in population size (Jones et al. 2021). Their survey methods, data quality and statistical methods were applied to a population that was isolated on an island with few predators (“lack of mammalian predators”, Langtimm et al. 1996) and thus not representative of typical mainland populations. That island population has since undergone a catastrophic decline due to fire and predation (Jones et al. 2021).

In this paper, we analyze a long-term record (1995-2023) of box turtle encounters from Jug Bay Wetlands Sanctuary, Lothian, Maryland using a Bayesian formulation of Jolly-Seber models (Royle and Dorazio 2008, ch. 10; Kery and Schaub 2010 ch. 10) to estimate annual population size, survival probability and recruitment rate. The Jug Bay data represent consistent sampling of the same population, producing sufficient sample sizes to obtain annual population size estimates as well as survival and recruitment. Our model uses encounters of individuals obtained by a combination of organized search effort and opportunistic encounters by us, Sanctuary staff, students and volunteers (“visual” encounters) and it includes information derived from individuals “found dead” using both telemetry and random encounters. Finally, the model includes individuals of unknown sex.

## STUDY AREA, METHODS, AND DATA ANALYSIS

The Jug Bay Wetlands Sanctuary covers about 688 ha (1,700 acres) on the Coastal Plain in Anne Arundel County, Maryland, along the eastern edge of the Patuxent River (Fig. 1). The Sanctuary is one of eight other large parks and nature reserves that protect over 4,000 ha in this section of the Patuxent River (Fig. 1). The study area in the Sanctuary was 100 ha. The diverse habitat consists of extensive freshwater tidal wetlands, mixed upland deciduous forest, riparian forest, stream valleys, managed meadows and adjacent agricultural fields. The area is drained by three semi-permanent streams which flow into the river including the respective floodplains. The “core study area” within which box turtle capture-recapture efforts were focused is 100 ha in size (Figure 1) and is based on a UTM grid which is designated on the ground by a series of 1-m tall PVC pipes. Turtles were encountered during regular visual searches and opportunistically by staff and visitors, we combine all such encounters together and refer to them as “visual encounters”. Quinlan and Swarth initiated this study and supervised the field work and surveys from 1995 to 2012; Quinlan supervised the field work from 2012 to 2023. At the time of publication, the survey data were not publicly available from the Jug Bay Wetlands Sanctuary.

**Figure 1.**
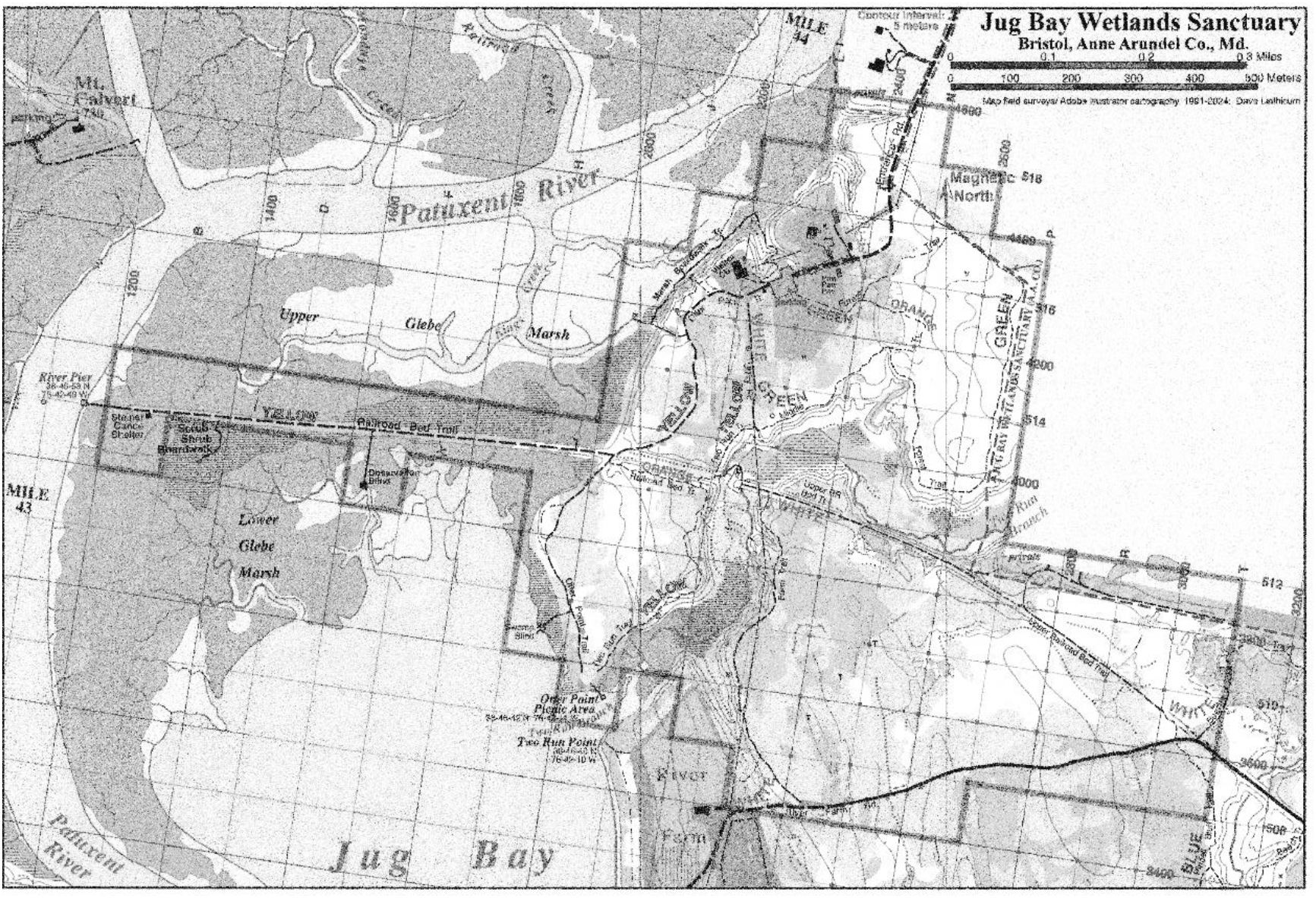
Jug Bay Wetlands Sanctuary core study area (thick polygon outline). (Map created by Jug Bay Wetlands Sanctuary staff)

### Lab and Field Methods

Turtles were captured in the study area and brought to the Wetland Center lab or processed in the field, where they were inspected, marked (after, Cagle 1939), measured, weighed, and sexed. One triangular notch was filed into two to four front and rear marginal scutes, giving each turtle a unique code. A photocopy or photographic image was taken of the plastron and placed in the record books. These two identification techniques made it possible to confirm the identify of each turtle when it was recaptured. Upon recapture, the scute notches were inspected and some ‘touched up’ with the file if necessary to make them more prominent. The straight-line carapace length and plastron width (at hinge) was measured to the nearest millimeter with pair of Haglof Mantax Aluminum calipers. Weight to the gram was taken on an Ohaus electronic scale. Sex was determined by assessing several secondary sex characteristics: presence or absence of a plastron depression, eye color, distance of cloaca from rear edge of carapace, and shape of the hind claws. After processing, each turtle was returned to the location where it was captured.

We discarded 365 encounter records of individuals not in the “core study area.” We discarded 18 encounters of individuals < 80 mm carapace length. This produced a data set containing 7,655 individual encounters of which 4,581 are telemetry encounters and the remainder are encounters by means other than telemetry (referred to as “random” – the result of random, opportunistic, organized search or other methods). These subsetting rules produced encounter histories of 572 individuals captured 1,998 times by visual encounter. Of these individuals, 189 were female, 308 were male and 75 were unknown sex (including 26 recorded as juveniles). A total of 74 previously marked individuals were found dead during the 29-year study. This includes 3 individuals that were found dead during telemetry encounters, and 71 found dead by visual encounter. We include information from these “found dead” encounters. Telemetry encounters were not used in the construction of encounter histories because nearly all individuals captured by telemetry in any given year were also encountered by visual search. Telemetry encounters do not provide additional information about Jolly-Seber model parameters in that case.

Because searching was done regularly between April and November each year, individuals may have been captured multiple times within the same year. For purposes of fitting discrete time Jolly-Seber models, we reduced encounter frequencies to binary events, *y*_*i, t*_, where *y*_*i,t*_ = 1 represents a capture in a given year and *y*_*i, t*_ = 0 represents non-capture.

The number of unique individuals captured each year is shown in Fig. 2. This shows two interesting features: (1) a large increase in total counts over the first 5-6 years of the monitoring effort and (2) a large decrease in observed counts over the subsequent 20+years. The total count from around 120 to 40 in recent years, a decline of roughly 67%. Of course, underlying this pattern here is the fact that survey effort was variable over time, and the pattern depicted in this figure does not account for that. In particular, the large increase in number of turtles encountered over the first five years is probably a result of increasing effort as the study was initiated, although survey effort was not recorded. Because survey effort was not recorded, we use statistical models for capture-recapture data – they allow us to estimate a year-specific detection probability which accommodates the variation over time in the survey effort. The least-squares linear trend (solid line in Fig. 2) fitted to the observed count produces a slope of -3.0685 individuals per year (t = -4.82, p < 0.0001), which is about -2.7% per year relative to the y-intercept of 111.85.

**Figure 2.**
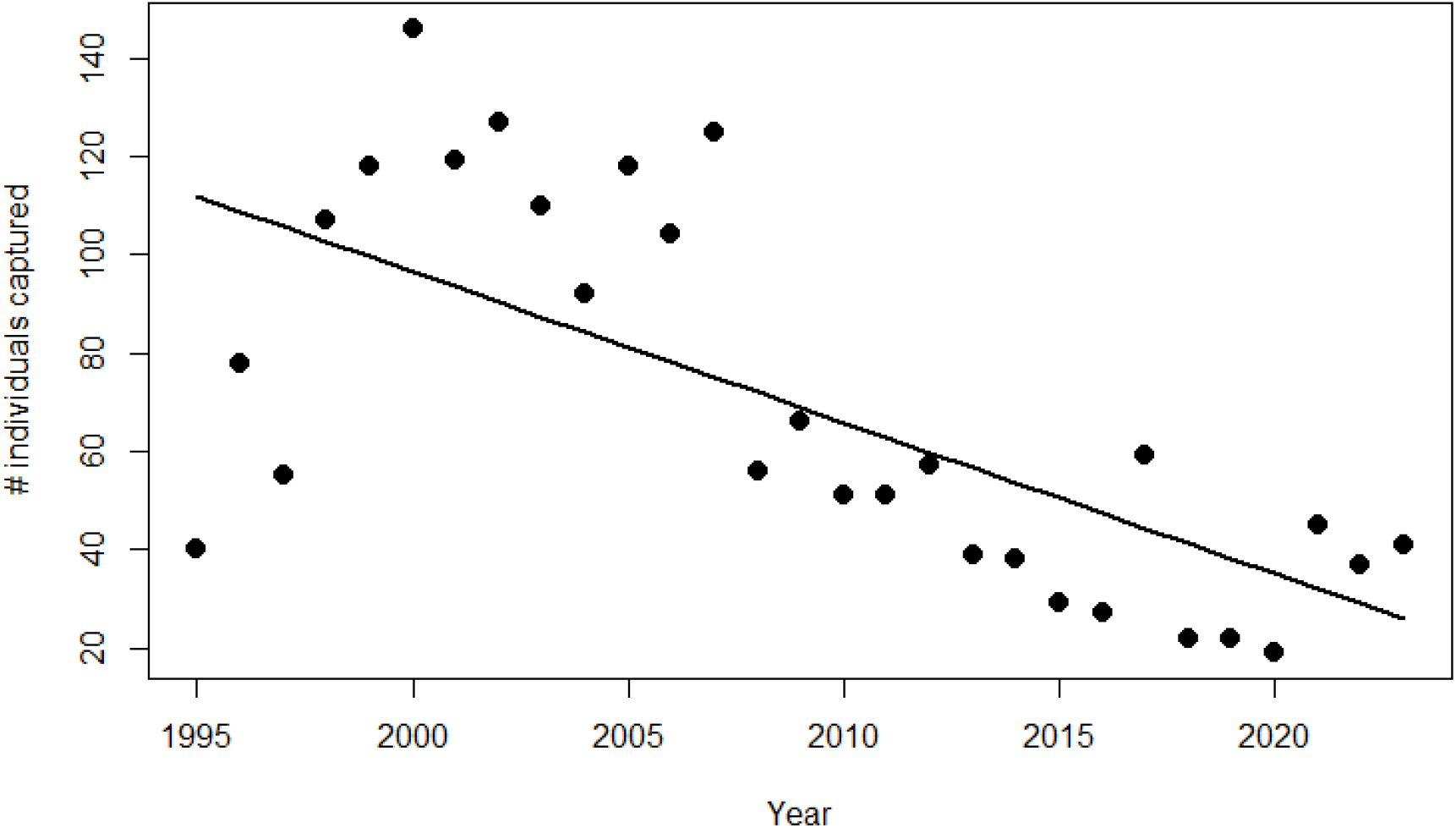
Number of unique individuals captured in each year. Solid line is the least-squares fit.

### The Jolly-Seber model

Joint estimation of population size and demographic parameters (survival, recruitment) from discrete-time encounter history data when individuals cannot be detected perfectly is usually done using the Jolly-Seber (JS) model (Schwarz 2001). We use a Bayesian hierarchical formulation of the model (Royle and Dorazio 2008) using an R-based (R Core Team 2019) code template from Kery and Schaub (2010) in which the model is fit in the JAGS software (Plummer 2003) using the jagsUI R package (Kellner 2015). This hierarchical or state-space formulation of the model (Royle 2008) describes individual survival, recruitment and detection of individuals in terms of distinct sub-models for the binary encounter observations *y*_*i,t*_ and the individual binary state variables *z*_*i,t*_ representing the “status” (alive or not alive) of each individual during each year or primary occasion.

A key idea of the Bayesian hierarchical formulation of the JS model that allows it to be conveniently fit in popular Bayesian analysis software such as JAGS is the use of data augmentation (Royle and Dorazio 2008) in which the observed encounter histories are embedded in a larger super-population of individuals including those that were not captured. Let *M* denote the size of this super-population. For each individual in the super-population that was not captured, the data are implied to be all-zero encounter histories, *y*_*i,t*_ = 0 for all *t* and then the model is defined for all *i* = 1, 2,…, *M* as follows. The encounter observations are assumed to be independent Bernoulli trials, conditional on the latent alive states, according to

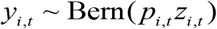

Where *p*_*i,t*_ is the probability of detecting individual *i* in year (or primary occasion) *t*, which may depend on individual- or year-specific covariates. Naturally, if an individual is not alive at time *t*, so that *z*_*i,t*_ = 0, then the observation *y*_*i, t*_ = 0 with probability 1 i.e., it is a deterministic 0. The alive states are binary (*z* =1 for alive, *z* = 0 for not alive) and assumed to be Markovian for *t* = 2,…,*T* according to

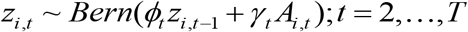

And, for *t* =1, *z*_*i*,1_ ∼ *Bern*(*ψ*_1_), where *ψ*_1_ represents the fraction of the super-population of individuals that are alive at time *t* =1. The variable *A*_*i, t*_ is a deterministic indicator of whether an individual in the super-population is available to be recruited just prior to primary occasion *t*, and it is a function of whether an individual has ever been alive whereas the parameters *γ* _*t*_ are pseudo-recruitment parameters, representing the fraction of individuals available to be recruited that are recruited in year *t*. This can be converted directly to a per-capita recruitment by tabulating the number of recruits each year as a step in the Markov chain Monte Carlo (MCMC) analysis and standardizing by the population size.

In general, individual and occasion-specific covariates can be modeled on both *p*_*i,t*_ and *ϕ*_*i,t*_. We assumed an additive “year + sex” model for detection probability and survival probability so that: *logit*(*p*_*i,t*_) = *α*_0_ *I* (*sex*_*i*_ = *male*) + *α*_*t*_ and *logit*(*ϕ*_*i,t*_) = *β*_0_ *I* (*sex*_*i*_ = *male*) + *β*_*t*_. Thus, *α* _0_ and *β* _0_ are the “male effect” on detection and survival, respectively. In addition, we assumed that *α*_*t*_ (female detection probability, on the logit-scale) were random effects, having a normal distribution: *α*_*t*_ ∼ Normal(*μ*_*α*_, *σ*_*α*_). We could have chosen to model the yearly survival parameters as random effects also, but we believe the data can support estimation of yearly survival as fixed effects and believe there is some biological interest in having “pure” estimates of survival not overly influenced by superfluous model structure.

The pseudo-recruitment parameters were assumed to vary by year as fixed effects. Because sex was missing for 75 individuals (26 Juveniles and 49 unknown adults), we assumed a prior distribution for the sex variable: *sex*_*i*_ ∼ Bern(*ψ* _*sex*_). We coded the variable *sex* as a binary variable with *sex* =1 representing a male and *sex* = 0 a female. Unknowns were coded in the data using the standard R representation of missing values *sex* = *NA*.

The JS model requires constraints on some of the detection parameters (Kery and Schaub 2010) which are not estimable under the full model with time-specificity of parameters. One common approach is to assume that the first and last detection probabilities are equal to their neighbors, i.e., p[1995] = p[1996] and p[2023] = p[2022]. This might be adequate if we could assume effort was approximately constant among years or an explicit effort covariate was measured and used as a covariate on *p*. However, we think effort was not approximately constant, and no covariate is available. Therefore, instead, we assumed *p* is year specific to accommodate variations in total effort over time and induced a weak stochastic constraint that the values of *p* are random effects from a common distribution by assuming that the logit-transformed yearly detection probabilities are normally distributed with mean *μ*_*p*_ and standard deviation *σ*_*p*_. This allows the first and last detection parameters to be estimated.

Several other modifications to the basic model were made. We regarded the visual encounters of turtles by all causes as stochastic events consistent with the assumptions of the JS model. In addition, during the course of the study 74 individuals were found dead, 3 of these were found by telemetry. Individuals found dead, regardless of cause, were regarded as deterministic encounters and only contribute information to the latent *z* states of the model. Specifically, during the year of dead encounter, subsequent z states are fixed at 0. As such, if an individual is found dead at some time say *t* then *z*_*i,t* +1_ and all subsequent states are set to 0, so that mortality occurred either in year *t* or some previous year. Visual encounter of dead individuals could be treated slightly differently – they could be assumed to be stochastic events with detection probability *p*, but in some cases the timing of death is highly uncertain so we just used them to fix the subsequent *z* states to 0 instead of trying to further refine the year of death.

We fit the model in the R package jagsUI (Kellner 2015) which is an R wrapper for the JAGS software (Plummer 2009). JAGS samples the posterior distribution of model parameters using MCMC and parameter estimates and their uncertainty are characterized by Monte Carlo averages of the posterior samples. Posterior summaries were based on 8 Markov chains run for 5000 iterations after a 1000 sample burn-in, with a thinning rate of 2, producing 20,000 total posterior samples. Convergence was assessed using the Gelman-Rubin “Rhat” statistic (Gelman et al. 2004). Values near 1 indicate convergence and, in practice, values < 1.1 are usually regarded as satisfactory. See Royle and Dorazio (2008) and Kery and Schaub (2010) for details on Bayesian analysis, MCMC, and the JS model.

## RESULTS

### Population size and trend

The estimated annual population size of box turtles each year from 1995 through 2023 is shown in Fig. 3 along with the upper and lower bounds of the 95% Bayesian credible interval of population size, N, for each year. The estimated population size trajectory shows a marked decline from the range N = 300 to N = 350 in the early years to close to N = 100 individuals by 2023. Using levels of N = 327 and N = 107 for the sake of calculation, the population decline is about 67% over the 29-year period. We computed the posterior distribution of the least-squares fit for a linear trend through the annual population sizes, producing the solid line in Fig. 3, and the posterior of the slope expressed relative to the intercept is shown in Fig. 4, indicating about negative 2.3% per year decline in population size. The posterior interval does not include 0. Thus, there is evidence of a long-term decline in the population size of box turtles at Jug Bay. We note that the population size being estimated by the model includes all individuals that might ever be available for capture on the 100-ha study area. This area is therefore larger than the nominal 100 ha study area, but the effective sample area is not well-defined absent models that account for spatial sampling (e.g., spatial capture-recapture; Royle et al. 2013).

**Figure 3.**
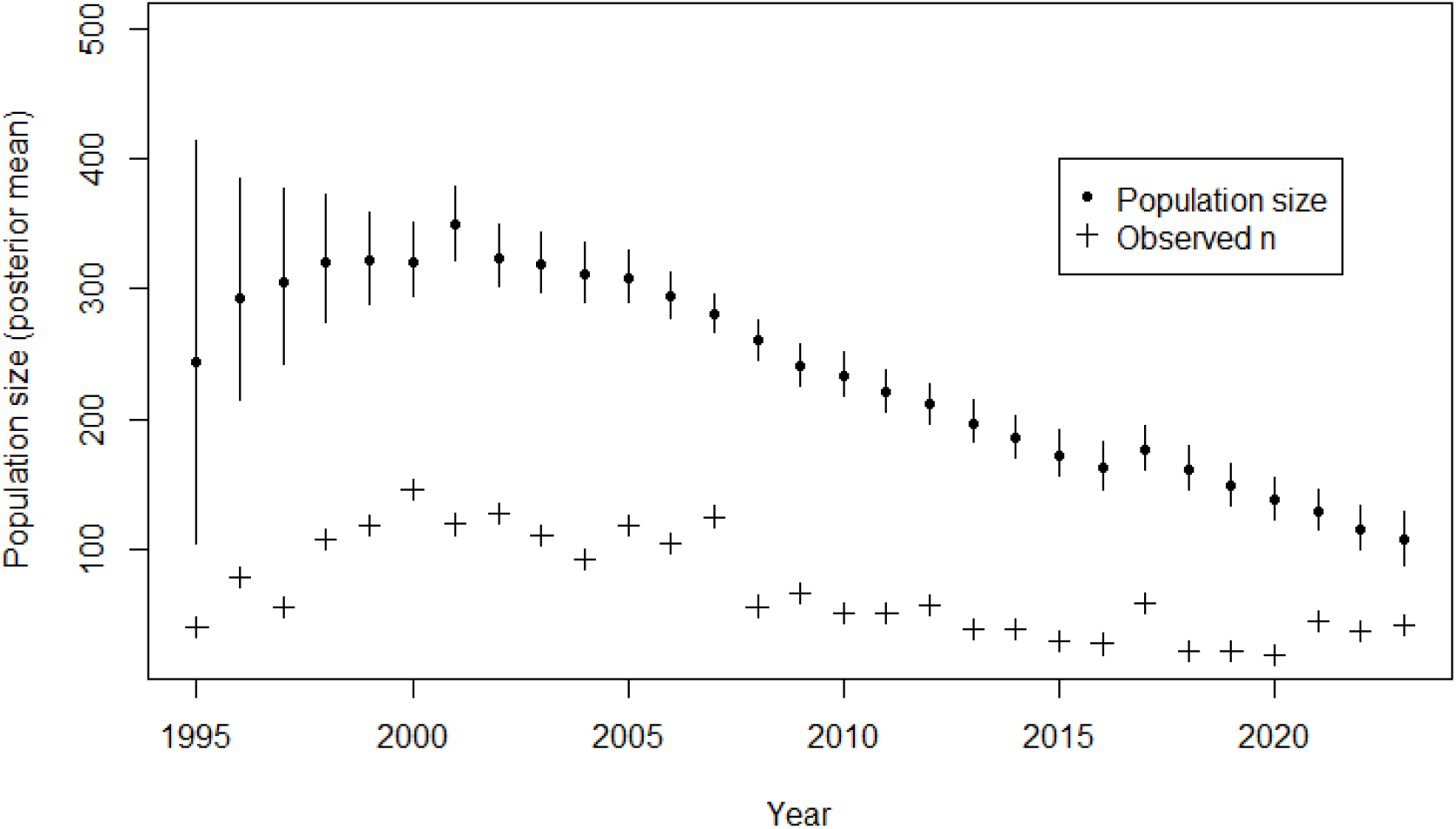
Estimated population size (black dots) and 95% posterior intervals (vertical lines). The number of unique individuals captured in each year is shown by the plus signs.

**Figure 4.**
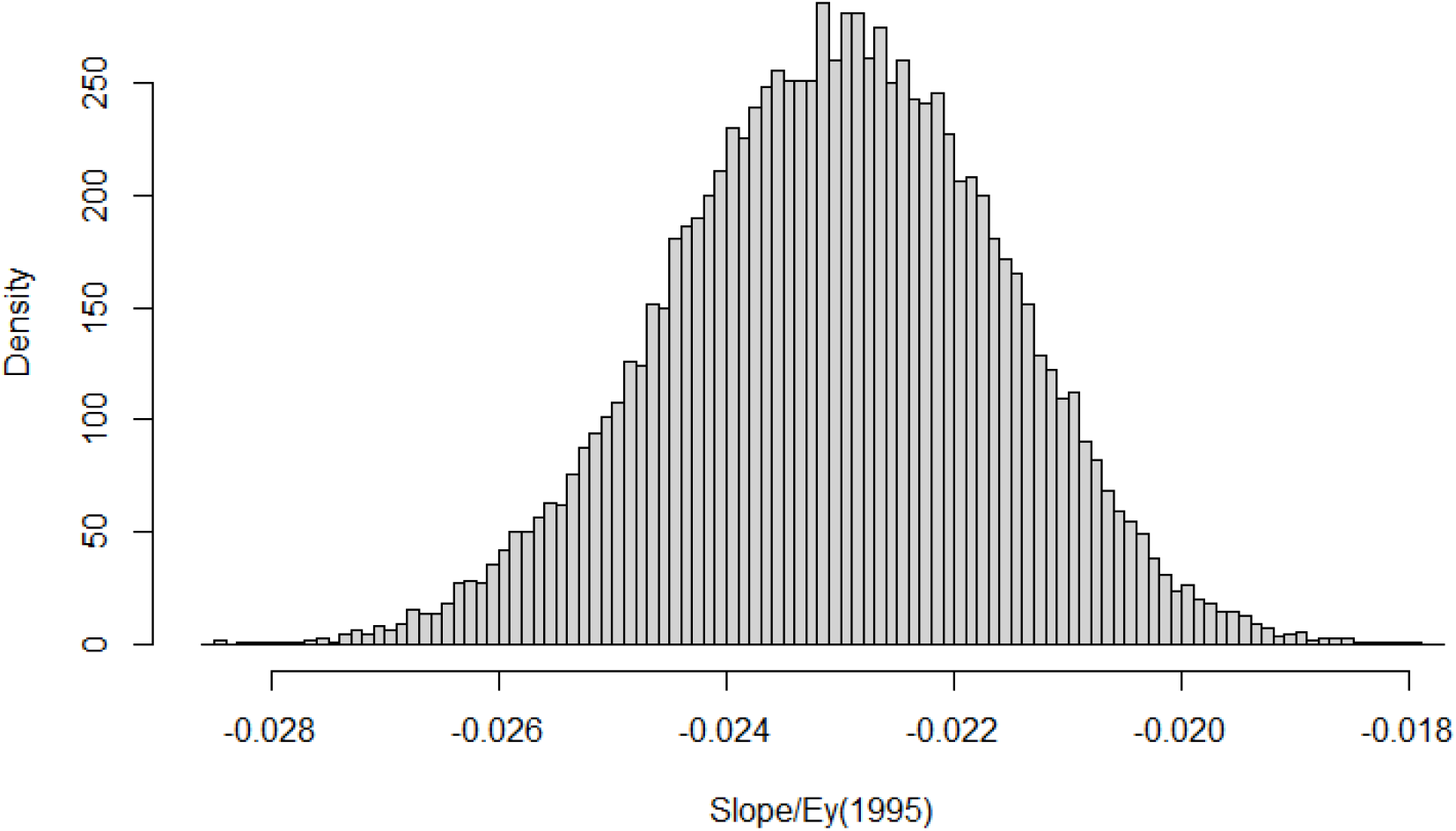
Posterior distribution of the linear rate of change in population size, defined as the slope of the least-squares fit to the posterior sample of population sizes *N*_*t*_ the expected population size in 1995.

### Model structural parameters

The main time-constant scalar parameters of the model are summarized in Table 1. The parameter *ψ* is the data augmentation parameter which we are not interested in interpreting directly, although the posterior mass should be away from 1.0 in order to ensure that estimates of *N* are not biased (Royle et al. 2013). The detection probabilities were assumed to follow a normal distribution on the logit-scale, and *μ*_*p*_ and *σ*_*p*_ are the mean and standard deviation parameters of that normal distribution. The posterior means of parameters *p*_*t*_ are shown in Fig. 5, but not summarized in Table 1. The parameters male.surv and male.p are the effect of ‘being male’ on survival and detection, respectively. We see a very large positive effect for males on survival (they survive at a higher rate), and a slightly negative effect on detection probability. Annual survival probabilities (posterior means and 95% credible intervals) for males and females are shown in Fig. 6. The parameter *ψ*_*sex*_ is the probability that an individual is male at the time of recruitment into the marked population. This estimate of 0.524 implies about 1.1 males per female at the time individuals enter the surveyed population (80 mm carapace length by our data sub-setting convention). Conversely, the sample proportion of males is 0.62 (308 males vs. 189 females in the sample), which is consistent with typical observed sex ratios. This favors males more because they are both slightly less detectable (male.p is negative) and survive at a much higher rate (male.surv is positive).

**Table 1.**
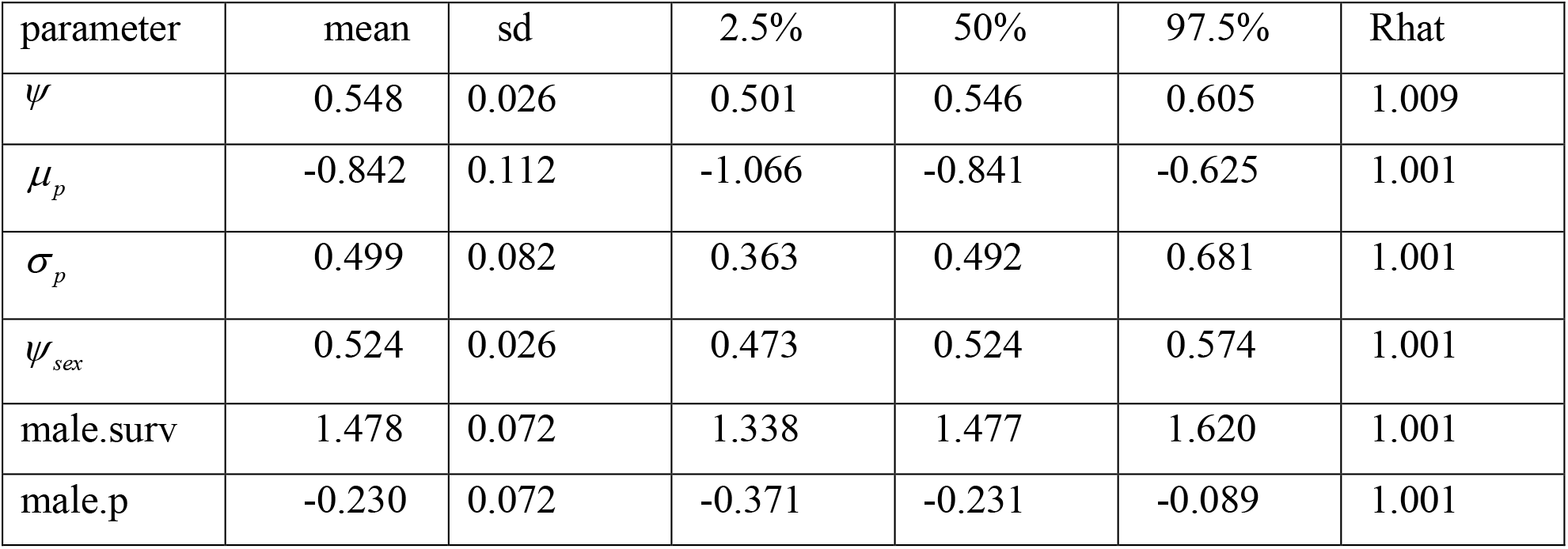
Posterior summaries (mean, standard deviation, specified percentiles) of the non-time-varying model structural parameters. *ψ* is the data augmentation parameter, *μ*_*p*_ is the mean of the logit-detection probabilities, *σ*_*p*_ is the standard deviation, *ψ*_*sex*_ is the population probability that an individual is a male, male.surv is the effect of being male on survival probability, and male.p is the effect of being male on detection probability. Rhat is the Gelman-Rubin convergence diagnostic.

**Figure 5.**
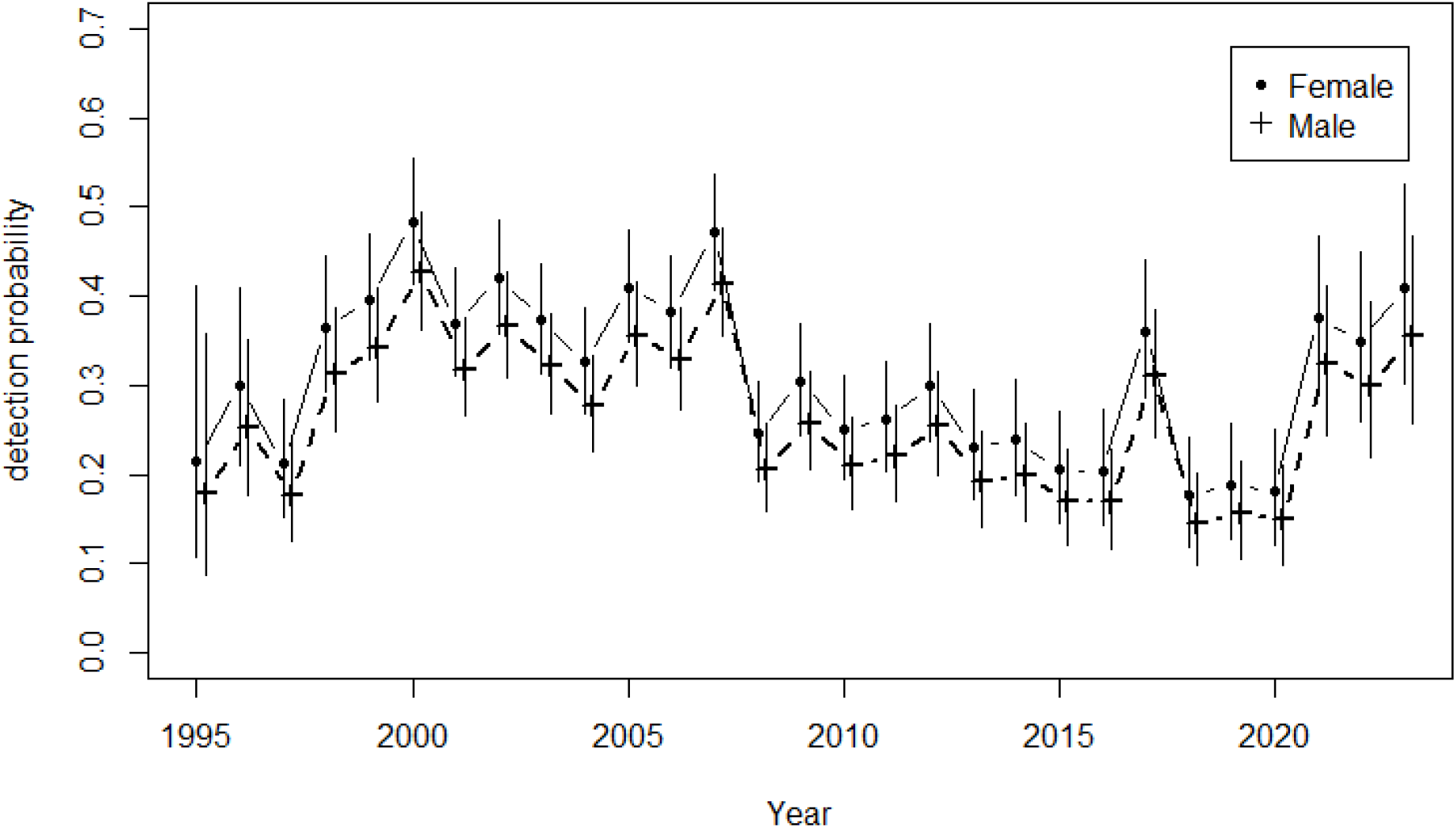
Estimated detection probability of box turtles at Jug Bay Wetlands Sanctuary for each year 1995-2023 with 95% Bayesian credible intervals (vertical lines) for females (black dot symbols) and males (plus signs). Male and female trajectories have a slight horizontal offset for visual clarity.

**Figure 6.**
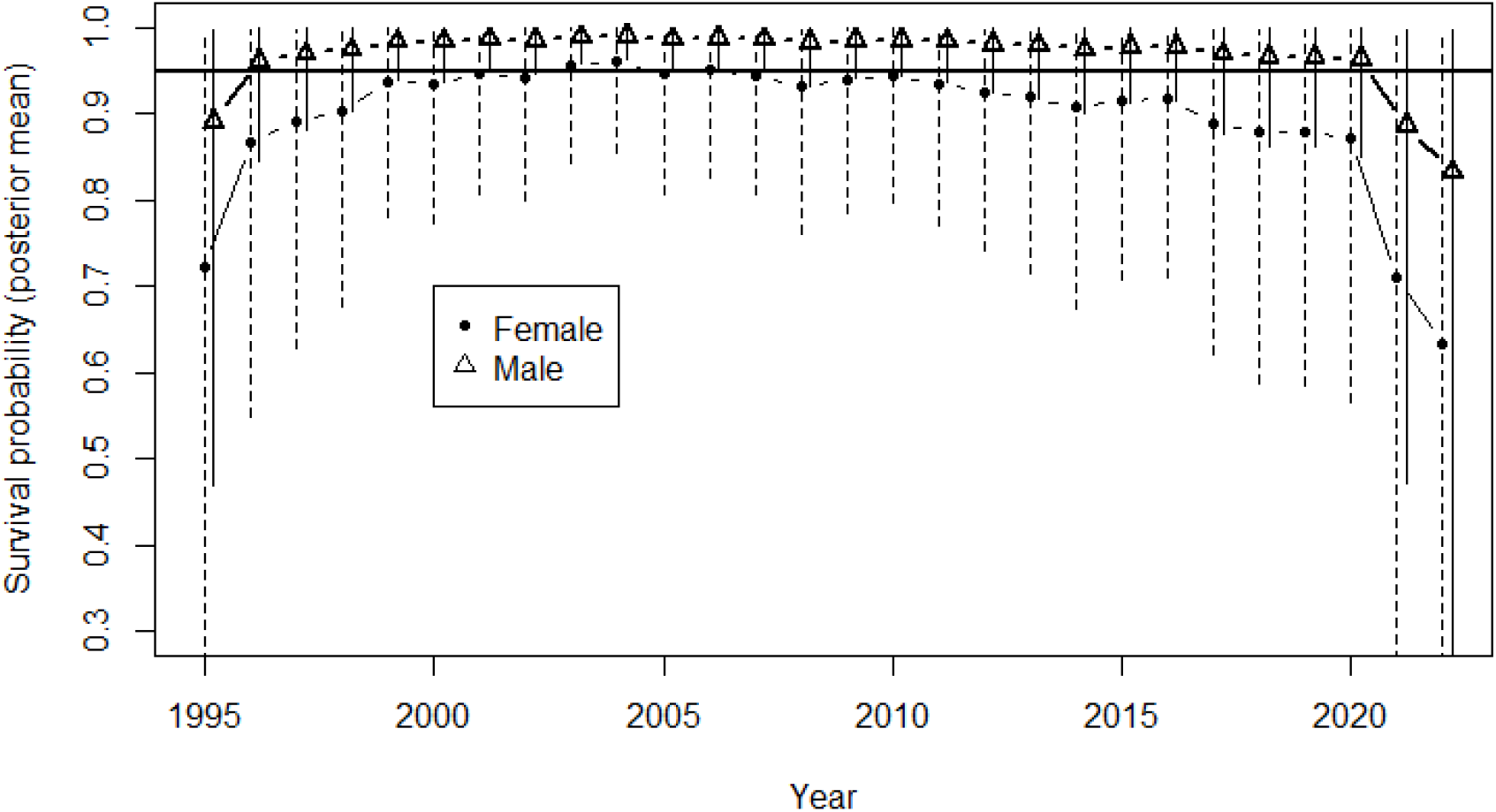
Estimated annual survival probability 1995-2023 of box turtles at Jug Bay Wetlands Sanctuary for males (triangles) and females (solid dots) with 95% Bayesian credible intervals (vertical lines). Horizontal black line is the value *ϕ* = 0.95.

### Capture probability

The estimated annual capture probability (posterior means and 95% credible intervals) for males and females over the 29-year period is shown in Fig. 5. An increase in the early part of the effort is evident, but generally annual capture probability fluctuates between 0.15 and 0.35.

### Survival and recruitment

Figure 6 annual survival for males and females. We note low precision at the beginning and end of the series, due to low sample sizes, and improved precision for interior years which is a consequence of the Markovian structure of the model and the “borrowing” of information that the model accomplishes by keeping track of individual identity over time. Data indicate that male survival is much higher than that of females (see Table 1 also). Fig. 7 shows the posterior distribution of average survival for males and females over the 29 period 1995 – 2023. Female average survival is 0.896 (95% CI: 0.855, 0.933) and male average survival is 0.968 (95% CI: 0.942, 0.983). Per-capita recruitment was estimated from the model as a derived parameter, by summing up the number of recruits at time *t* (individuals that entered the modeled-population at time *t* but were not alive at time *t* −1) and dividing by the population size at time *t* − _1. We further averaged these annual estimates over 4-year periods to reduce variation. The posterior distributions are shown as box plots in Fig. 8 (left). The observed number of individuals captured having carapace length between 80 and 120 mm is shown in the right panel of Fig. 8.

**Figure 7.**
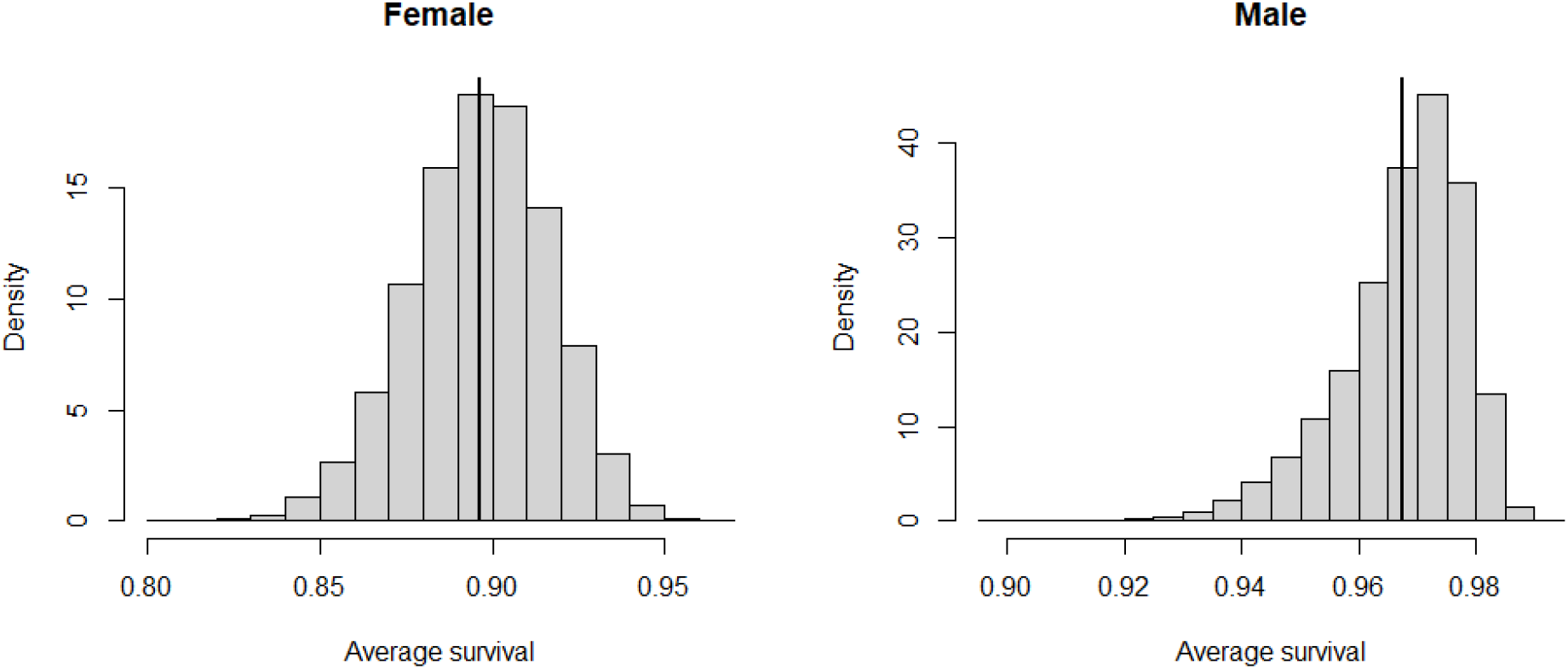
Average survival of Eastern box turtles over the period 1995-2023 at Jug Bay Wetlands Sanctuary for females (left) and males (right). Horizontal line is the value *ϕ* = 0.95.

**Figure 8.**
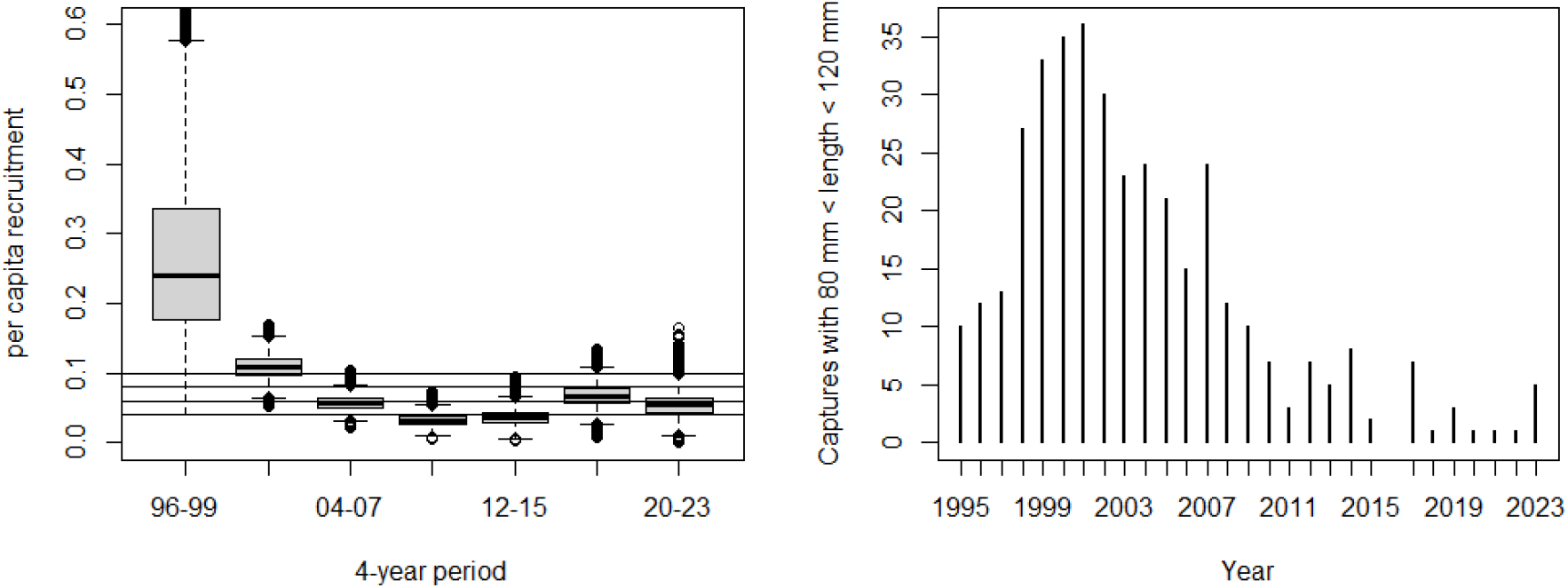
Left: Posterior distributions (shown as box plots) of average per capita recruitment per 4-year period under the Jolly-Seber model. Estimates were averaged over 4-year periods to reduce variation. Horizontal lines mark values of 0.10, 0.09, 0.08, and 0.07 recruits per adult individual. Right: Observed number of captures of individuals with carapace length > 80 mm and < 120 mm.

## DISCUSSION

We analyzed a nearly three-decade long capture-recapture data set on Eastern box turtles from a protected site on the Coastal Plain of Maryland, in order to estimate annual population size, trend, and demographic parameters (survival and recruitment). The estimated decline in box turtle population size that we found, about 67% over 29 years in the Jug Bay Wetlands Sanctuary, is noteworthy because, to the best of our knowledge, our time-series of population size estimates and survival probabilities are the longest and most-complete for any population of Eastern box turtles. Our observed population decline of about 67% is consistent with other long-term trend estimates: about 80% over 50 years reported by Hall et al. (1999), 75% over 42 years by Kemp et al. (2022) and 76% over 34 years Nazdrowicz et al. (2008). These long-term estimates are remarkably consistent and, coincidentally, all represent populations in close proximity to one another in the mid-Atlantic region (Maryland, Pennsylvania and Delaware, respectively). Our analysis of monitoring data from a Maryland site different than that of Hall et al. (1999), taken together with these other trend estimates, provides strong evidence of systematic declines in Eastern box turtle populations in the mid-Atlantic region. Furthermore, these observed long-term declines are not small numbers. These populations have declined rapidly and, at least for the population we considered in this paper, continue to decline (Fig. 3).

Over the 29-year period we estimated that female survival averaged 0.896 (95% CI: 0.855, 0.933) and male survival averaged 0.968 (95% CI: 0.942, 0.983). There have been many estimates of Eastern box turtle survival reported in the literature, from both telemetry and capture-recapture studies. Most studies report “overall” survival which is constant over time and sometimes averaged over sexes, so our results may not be directly comparable to many published estimates, although they are reasonably consistent in magnitude for both sexes for comparable reports. Roe et al. (2021) reported 90.7 - 96.8 survival depending on the demographic group, and lower survival for females consistent with ours. Nazdrowicz et al. (2008) reported an average survival (both sexes) of 0.813, 0.945, 0.951, 0.977 for four small populations; Currylow et al. (2011) reported overall survival of 0.962 for all adults; Brisbin et al. (2008) overall apparent survival of 0.954 (males and females). Our observed sex difference (higher for males) is typical for the species and consistent with increased mortality due to nesting activity and larger space use of females (Habeck et al. 2019; Bulte et al in prep), and also owing to seasonal movements to and from suitable nesting areas, which forces some females to cross roads. Our estimates of effective per-capita recruitment, defined as the number of individuals > 80 mm entering the population at time *t* divided by the population size at time *t* −1 show a steep decline from the start of monitoring in 1995 through the period 2008-2015 before rebounding slightly. The pattern is consistent with the observed frequency of capture of small-sized individuals between 80- and 120-mm carapace length. Conversely, there appears to be no systematic change in survival probability over the period although the beginning and end of the record show highly imprecise estimates which produce unstable point estimates of survival with very diffuse posterior distributions. This is because, in the JS model, there is relatively little information about survival until sufficient individuals have been marked and recaptured, and information about survival propagates both forward and backward in time due to the Markovian structure of the JS model. The pattern in recruitment, supported by the empirical counts of small-sized individuals, and the lack of a systematic decline in survival, suggests that the overall population decline over this period is likely due to reduced recruitment. The site has been protected from development and unrestricted access over the period of data collection and has contained a relatively stable habitat structure over this period. The causes for reduced recruitment are unclear and we are unable to provide insight into possible causes for reduced recruitment levels in our study population. Researchers have identified many factors or threats that impact Eastern box turtle populations (see Erb and Roberts 2023), but evidence for the relative importance of these factors is scant. Of the causes that might operate in our study site, predation on nests and young, and disease are possible. Over the course of this study, three marked turtles were discovered crushed by vehicles on dirt roads. Finally, the Sanctuary is open on a limited basis to the public, however, we have never encountered a visitor collecting a turtle. Confirmation of this pattern in recruitment for other populations could enhance our understanding of possible causes of reduced recruitment in the species.

Our estimates of survival and recruitment parameters could be useful in aiding the conservation of this species. For example, they could be used to train models for population status assessments or population viability analysis (PVA), e.g., see Moore et al. (2022). Additionally, consistent long-term data collection at other box turtle populations could strengthen the model. A challenge in monitoring populations of the Eastern box turtle is that monitoring has to be done over a very long time in order to detect trends and patterns in demographic parameters, due to the high survival and low recruitment of the species. While the Eastern box turtle may be one of the most monitored reptile species in North America, with many efforts initiated at local sanctuaries, schools and communities, few studies persist long enough to generate quality long-term data with sufficient sample sizes to estimate demographic parameters.

We applied the JS model to the long-term data from Jug Bay Wetlands Sanctuary. This model is used for inference about population demography from animal population studies in which individuals are detected imperfectly (capture probability < 1). Our results emphasize the importance of using statistical methods such as the JS model to interpret monitoring data. We were able to produce statistically qualified estimates of recruitment, sex-structured survival and population size over a period of record that, to the best of our knowledge, is unequaled for this species. Accounting for imperfect and sex-structured detection probability shows the expected ‘adjustment’ in sex ratio from near 1:1 “at recruitment” (80 mm carapace length) to strongly favoring males (about 1.6:1) in the adult population at large, which is typical in box turtle populations (Dodd 2002). Allowing for detection probability to vary over time is important for interpretation of changes in counts when environmental conditions and effort also vary over time, as is most likely the case in the Jug Bay data set where we note that annual detection probability shows an increase from the initiation of the effort in 1995 for about 6 years and then reaching low levels of detection again in 1998-2000. Our estimates of population size are “free” from biasing effects of this temporal variability in detection probability and thus interpreted directly as changes in actual population size.

Despite all the advantages of formal analysis using the JS model, a number of improvements in the data collection and analysis could be made. For example, we treated each year as a single sampling occasion and thus an individual was captured (y = 1) if it had 1 or more captures in that year. This disregards many recaptures (some individuals we recaptured dozens of times in a year), but it is difficult to accommodate the recapture information without having more precise and curated information about spatial and temporal effort of sampling. One might consider dividing a season up into smaller sub-samples within a year, this is called the “robust design” (Kendall and Pollock 1992). Although this may lead to more statistically efficient estimation, it would probably lead to additional problems in modeling temporal variation in detection probability because effort is likely to be more heterogeneous within a year. In addition, spatial information about search effort and capture locations could be used in spatially explicit capture-recapture models (Royle and Turner 2022), which could greatly increase the precision of detection probability estimates and hence demographic parameters. However, for the Jug Bay data set there is not consistent information about search effort as many of the encounters are opportunistic. In general, we recommend that monitoring efforts for turtles should attempt to record survey effort as much as possible, including both spatial and temporal effort.

## ACKNOWLEDGEMENTS

The following individuals devoted many days in the field searching for, capturing, identifying, measuring, and marking box turtles: Susan Blackstone, Bob Williams, Sandy Teliak, Sandy Barnett, Les Silva, Susan Hagood, Terry Duckett, Susan Matthews, Michael Marchand, Antonio Cordero, Karyn Molines, Elaine Friebele, Lindsay Hollister, Liana Vitali, Kathy Chow, Siobhan Percey, Mary Kay Sistik, Allison Burnett, Eva Blockstein, Gregory Bulte, Morgan Angus, Melissa Bennett, Krista Capps, Josh Capps, Susan Curless, Brett DeGregorio, Amber Heramb, Anna Moyer, Jennifer Lentz, Ben Mattics, Beth Nicholls, Logan Olds, Joe Sage, Gavin Studds, Ramona Sampsell, Evan Swarth, Lisa Thurston, Tina Whittle, Tara Whittle, Katy Clark, Sam Kuo, Michelle Lacombe, Jen Zimmerman, Jeanette Kazmierczak, Travis and Jessica Roney, Michelle Campbell, Christina Olson, Max Maddox, Felicity Kreger, Jamie Parker, Alison Kaufman, Lynette Fullerton, Charlotte Weinstein, Kevin Creek, Madison Sanders, Colleen McCluskey, and Katherine Baer. We thank David Linthicum for creating the study area map and Clint Cosner for surveying the grid pole locations. We especially thank Sanctuary Director Dr. Patricia Delgado for her on-going support. Funding for this study was provided by the Anne Arundel County Department of Recreation and Parks, the Friends of Jug Bay, and the Chesapeake Bay National Estuarine Research Reserve. We thank Lori Erb for reviewing a draft of the manuscript. Any use of trade, firm, or product names is for descriptive purposes only and does not imply endorsement by the U.S. Government. Our study was carried out under Collecting Permit #55329, Maryland Department of Natural Resources.

